# Comparative analysis of single-cell and single-nucleus RNA-sequencing in a rabbit model of retinal detachment-related proliferative vitreoretinopathy

**DOI:** 10.1101/2022.11.07.515504

**Authors:** Clayton P. Santiago, Megan Y. Gimmen, Yuchen Lu, Minda M. McNally, Leighton H. Duncan, Tyler J. Creamer, Linda D. Orzolek, Seth Blackshaw, Mandeep S. Singh

## Abstract

**Purpose:** Proliferative vitreoretinopathy (PVR) is the most common cause of failure of retinal reattachment surgery and the molecular changes leading to this aberrant wound healing process is currently unknown. We aimed to study PVR pathogenesis using single-cell transcriptomics to dissect cellular heterogeneity in a rabbit PVR model.

**Methods:** PVR was induced unilaterally in Dutch Belted rabbits. At different timepoints following PVR induction, retinas were dissociated into either cells or nuclei suspension and processed for single-cell or single-nucleus RNA sequencing (scRNA-seq or snRNA-seq).

**Results:** scRNA-Seq and snRNA-Seq were conducted on retinas at 4 hours and 14 days after disease induction. While the capture rate of unique molecular identifiers (UMI) and genes were greater in scRNA-seq samples, overall gene expression profiles of individual cell types were highly correlated between scRNA-seq and snRNA-seq. A major disparity between the two sequencing modalities is the cell type capture rate, however, with glial cell types over-represented in scRNA-seq, and inner retinal neurons were enriched by snRNA-seq. Furthermore, fibrotic Müller glia were over-represented in snRNA-seq samples, while reactive Müller glia were in scRNA-seq samples. Trajectory analyses were similar between the two methods, allowing for the combined analysis of the scRNA-seq and snRNA-seq datasets.

**Conclusions:** These findings highlight limitations of both scRNA-seq and snRNA-seq analysis and imply that use of both techniques can more accurately identify transcriptional networks critical for aberrant fibrogenesis in PVR.

## Introduction

Proliferative vitreoretinopathy (PVR) is among the most important causes of visual morbidity in patients with retinal diseases, and its treatment is a major unmet need. PVR is analogous to an aberrant wound healing process that stiffens and distorts the normally pliant retina and disrupts its natural conformance to the curved posterior eye wall ^1^. Early-stage PVR is characterized by retinal stiffening and contraction and retinal pigment epithelium (RPE) cells and activated Müller glia are thought to play important roles at this stage ^2,3^. Disease progress involves retinal traction and detachment, and ultimately vision loss. Triggers of PVR include retinal detachment, infection or inflammation, penetrating injury, hemorrhage, and most frustratingly, the incisional retinal surgery that is often used to treat these conditions ^4^. Multiple episodes of reparative surgery are often required to reposition the retina and mitigate against further vision loss.

Aside from RPE cells and Müller glia, multiple other retinal cell types may be involved in early and/or late PVR development, yet the temporal cascade of molecular changes in various cell types remain poorly understood. This knowledge gap is further compounded by processes that cause cells to substantially alter their gene expression profiles, such as epithelial-to-mesenchymal transition in which RPE cells assume contractile properties^5^. In general, molecular data regarding PVR have come from bulk analysis ^6,7^. The lack of information on temporally-resolved cell-specific gene expression changes has impeded research on developing rational pharmacologic treatment to target the specific pathways that initiate or drive progression of PVR.

To address this knowledge gap, we aimed to study PVR pathogenesis using single-cell transcriptomic analysis. Over the last few years, this approach has been used in multiple species to identify molecular markers of virtually all retinal cell subtypes ^8–10^, identify gene regulatory networks controlling retinal development and regeneration ^11–14^, and identify molecular changes associated with onset and progression of disease ^15–17^. Single-cell transcriptomics can be conducted using single-cell RNA-Seq (scRNA-Seq), in which whole dissociated cells are profiled, or using single-nucleus RNA-Seq (snRNA-Seq) on isolated cell nuclei ^18,19^. Both approaches have been reported to have specific advantages and disadvantages ^20–22^. ScRNA-Seq captures larger numbers of fully-spliced mRNA species, capturing both cytoplasmic and nuclear transcripts, but is not usable in analysis of frozen or fixed tissue. SnRNA-Seq, on the other hand, is suitable for analysis of archival material and efficiently captures nuclear transcripts but is enriched for un-or partially-spliced transcripts ^23,24^. Selective loss of individual cell types has also been reported for the two approaches, with loss of large, mature neuronal subtypes in brain and retina, and fibrotic cells in the kidney with scRNA-Seq ^12,25–27^. In contrast, depletion of active microglia has been reported in brain snRNA-Seq libraries ^28^.

Though both scRNA-Seq and snRNA-Seq have been extensively used in the retina, a systematic comparison of the two approaches using the same input material has not been conducted. As a result, it is not clear to what extent the limitations of the two approaches apply, particularly in the study of complex retinal disease. Likewise, little is known about the relative accuracy and efficiency of these methods in the retina of adult mammalian species other than mice. In this study, we use both scRNA-Seq and snRNA-Seq to globally profile cellular transcriptomic changes in a rabbit surgical model of proliferative vitreoretinopathy (PVR). In addition to large differences between the two approaches in the relative abundance of major cell types, we also observe differences in the efficiency of capture of fibrotic and reactive Müller glia, differences in the overall levels of cell-cell contamination, and differential expression of many individual genes. These findings highlight the limitations of each approach and imply their combined use may prove more effective at profiling global transcriptional changes in disease.

## Results

To globally profile gene expression changes that occur during the progression of PVR, we conducted scRNA-Seq and snRNA-Seq on retina dissected from rabbits in which PVR-like lesions had been induced by vitrectomy, retinal detachment, induced by saline injection, platelet-enriched plasma injection, and cryotreatment to induce scar formation ^29,30^. Retina tissue was extracted from the lesion site, dissociated, split into two equal portions, and then processed and analyzed using scRNA-Seq or snRNA-Seq, respectively (Fig. 1A). Three different treatment conditions were profiled – uninjured control, and both 4 hours and 14 days following PVR induction – thereby ensuring a broad representation of disease progression.

**Figure 1:**
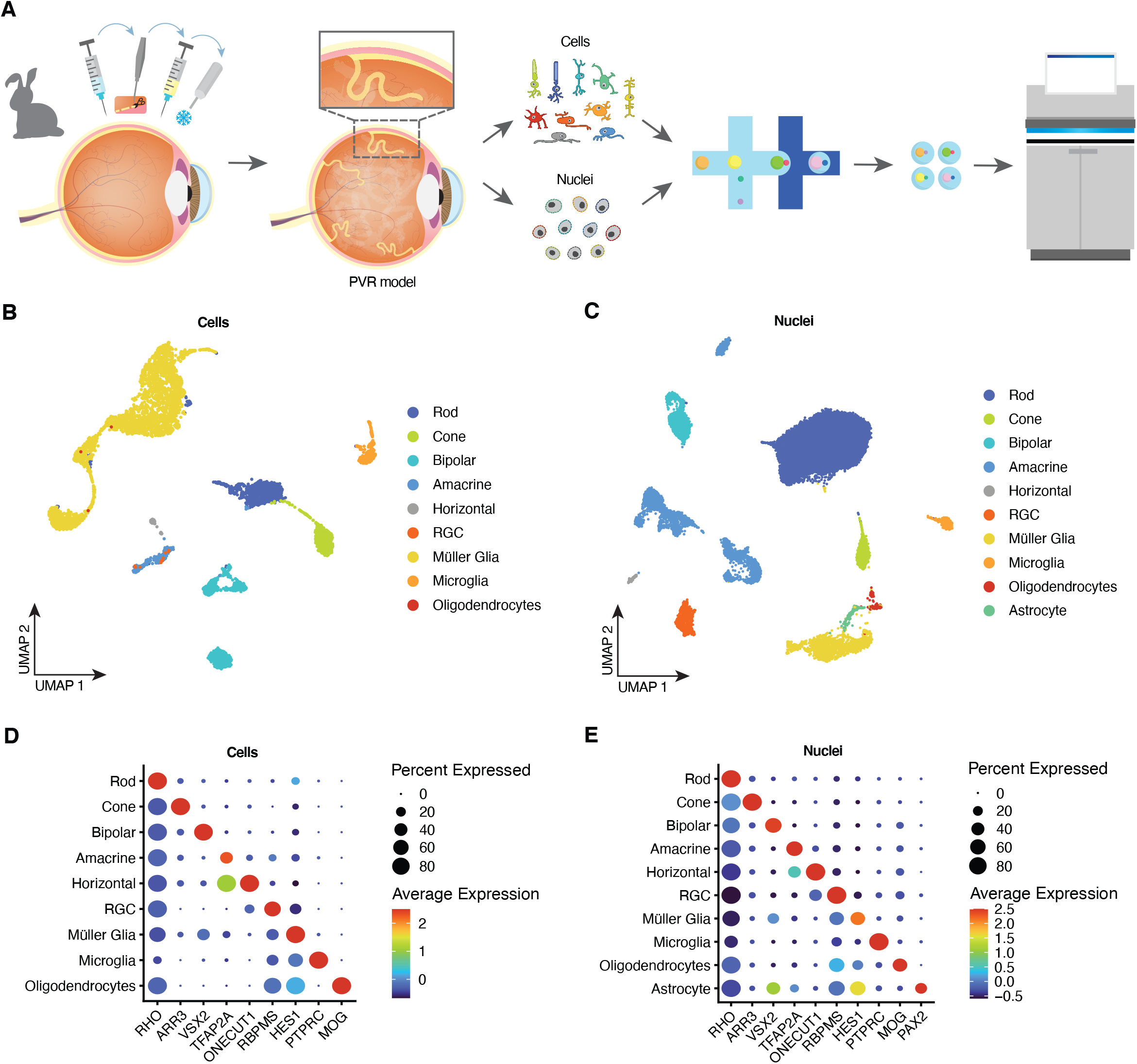
All major retinal cell types are captured by scRNA-Seq and snRNA-Seq. (A) Schematic summary of the study. PVR lesions were induced via lensectomy, pars plana vitrectomy, and retinotomy followed by retinal cryotherapy and intravitreal autologous platelet-rich plasma injection. At different time points, the lesions were dissected out and cells were dissociated, or nuclei were isolated before profiling by single cell or single nuclei RNA-seq. (B-C) Combined UMAP projection of all cells (left) or nuclei (right) profiled in this study. (D-E) Examples of gene expression levels for selected cell type-specific genes for cells (left) or nuclei (right).

We then combined datasets from all three timepoints for scRNA-Seq (Fig. 1B) and snRNA-Seq (Fig. 1C) separately, visualizing different cell types using UMAP analysis. Using well-characterized cell type-specific marker genes, we were able to readily identify every retina major cell type in each dataset, with the exception of astrocytes, which could be clearly resolved in the scRNA-Seq dataset (Fig. 1D,E, Supplementary Fig. S1A,B). Oligodendrocytes, which are not present in mouse or human retina, are abundant in the rabbit retina, and myelinate the axons of retinal ganglion cells in structures known as the medullary rays ^31^.

We next combined these two datasets and quantified the abundance of each major cell type. Overall gene expression profiles were very similar for individual cell types, and combined UMAP analysis revealed that common cell type-specific clusters detected by scRNA-Seq and snRNA-Seq were essentially superimposable in the combined UMAP plot (Fig. 2A). Direct comparison of cell type-specific gene expression profiles likewise showed very high correlation between scRNA-Seq and snRNA-Seq profiles of individual cell types, with these invariably being more closely correlated than profiles of other cell types (Fig. 2B).

**Figure 2:**
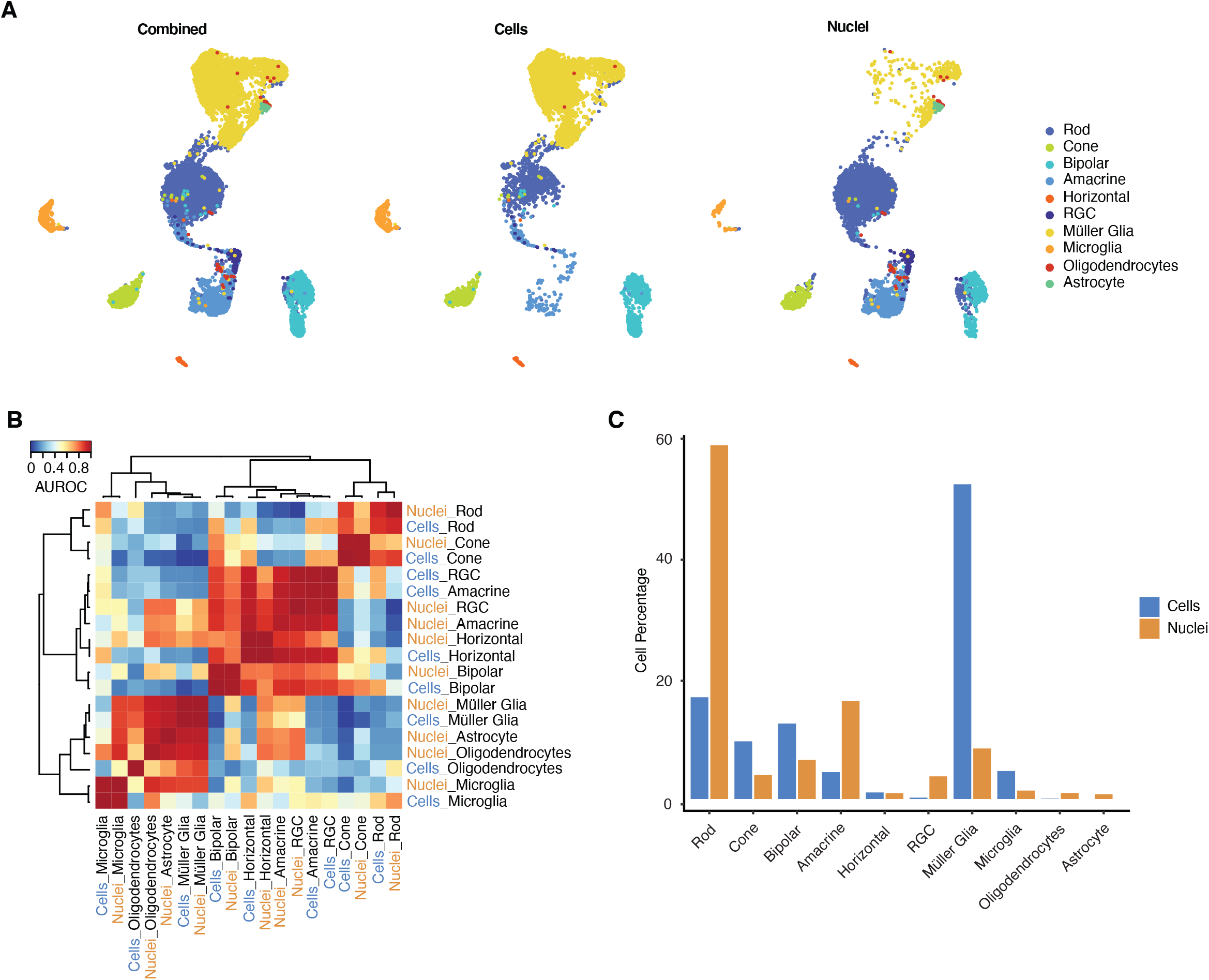
snRNA-seq has improved capture efficiencies for neuronal cell types. (A) UMAP embedding of scRNA-Seq and snRNA-Seq profiles showing profiles of the combined dataset (left) scRNA-Seq (middle) and snRNA-Seq (right), colored by the assigned cell type signatures. (B) Heatmap of Area Under the Receiver Operating Characteristics (AUROC) scores of the combined dataset between all retinal cell types based on highly variable genes. All replicating cell types cluster together after applying hierarchical clustering. (C) Bar plots of cell type proportions from each sequencing modality.

However, we observed major differences in the relative abundance of each cell type. Müller glia, cone photoreceptors, bipolar cells, and microglia were all overrepresented in the scRNA-Seq dataset, while rod photoreceptors, amacrine cells, retinal ganglion cells, and oligodendrocytes were enriched in the snRNA-Seq dataset. Horizontal cells were roughly equally abundant in both samples (Fig. 2C, Supplementary Fig. S2A). Though a comprehensive histological analysis of cell type ratios has yet to be conducted in rabbit retina, it is clear that the snRNA-Seq dataset provides an overall more accurately reflection of the true abundance of these various cell types. For instance, the rod:cone ratio as observed histologically is 20:1 ^32^, 13:1 as measured by snRNA-Seq and 1.7:1 as measured by scRNA-Seq (Supplementary Fig. S2C). Likewise, the histologically-defined relative ratios of inner retinal cells are 27 (bipolar):21 (amacrine):17 (Müller glia):1 (horizontal cells) ^33^. We observe that these are 12:4:48:1 for scRNA-Seq and 7:17:9:1 for snRNA-Seq (Fig. 2C, Supplementary Fig. S2B). Since roughly half of all amacrine cells exist as displaced amacrine cells in the ganglion cell layer ^34^, the ratios of cells profiled using snRNA-Seq closely match those defined using histological approaches.

We next more closely examine differences in the gene expression levels between the scRNA-Seq and snRNA-Seq datasets. As expected, when data from all individual cellular or nuclear profiles are aggregated, we observe significantly greater numbers of unique molecular identifiers (UMI) and individual genes detected in scRNA-Seq relative to snRNA-Seq profiles (Fig. 3A). We likewise observe that expression of individual genes are detected in a larger fraction of individual scRNA-Seq than snRNA-Seq profiles (Fig. 3B). As expected, we also observed a vastly larger number of spliced reads in the scRNA-Seq dataset when compared to the snRNA-seq dataset (Supplementary Fig. S3A). Correspondingly, we observed both common and cell type-specific differences in the expression levels of mRNAs corresponding to specific functional classes of genes in the scRNA-Seq relative to the snRNA-Seq dataset (Fig. 3C). Transcripts encoded by genes controlling protein synthesis, oxidative phosphorylation, viral gene expression, synapse formation, and RNA metabolism and splicing were enriched in scRNA-Seq samples from all cell types, while snRNA-Seq samples are enriched for transcripts encoding genes regulating histone modification, GTPase activity and cellular morphogenesis, and dendrite development. Notable cell type-specific differences in measured gene expression levels included enrichment of transcripts regulating protein dephosphorylation and ciliogenesis in cone snRNA-Seq, polyubiquitination in microglial snRNA-Seq, and neutrophil degranulation in microglial scRNA-Seq datasets. Overall, we observed 2,401 genes showing significantly higher expression in the snRNA-Seq dataset, while in the scRNA-Seq dataset we observed 2,093 to show higher expression in one or more cell types (Fig. 3D, Supplementary Fig. S3C-I). We also noted substantial differences in the level of contamination of individual cell types by genes specifically expressed in other cell types in the two datasets, with snRNA-Seq overall showing lower levels of contamination relative to scRNA-Seq data, particularly for rod photoreceptor-enriched genes (Supplementary Fig. S3B).

**Figure 3:**
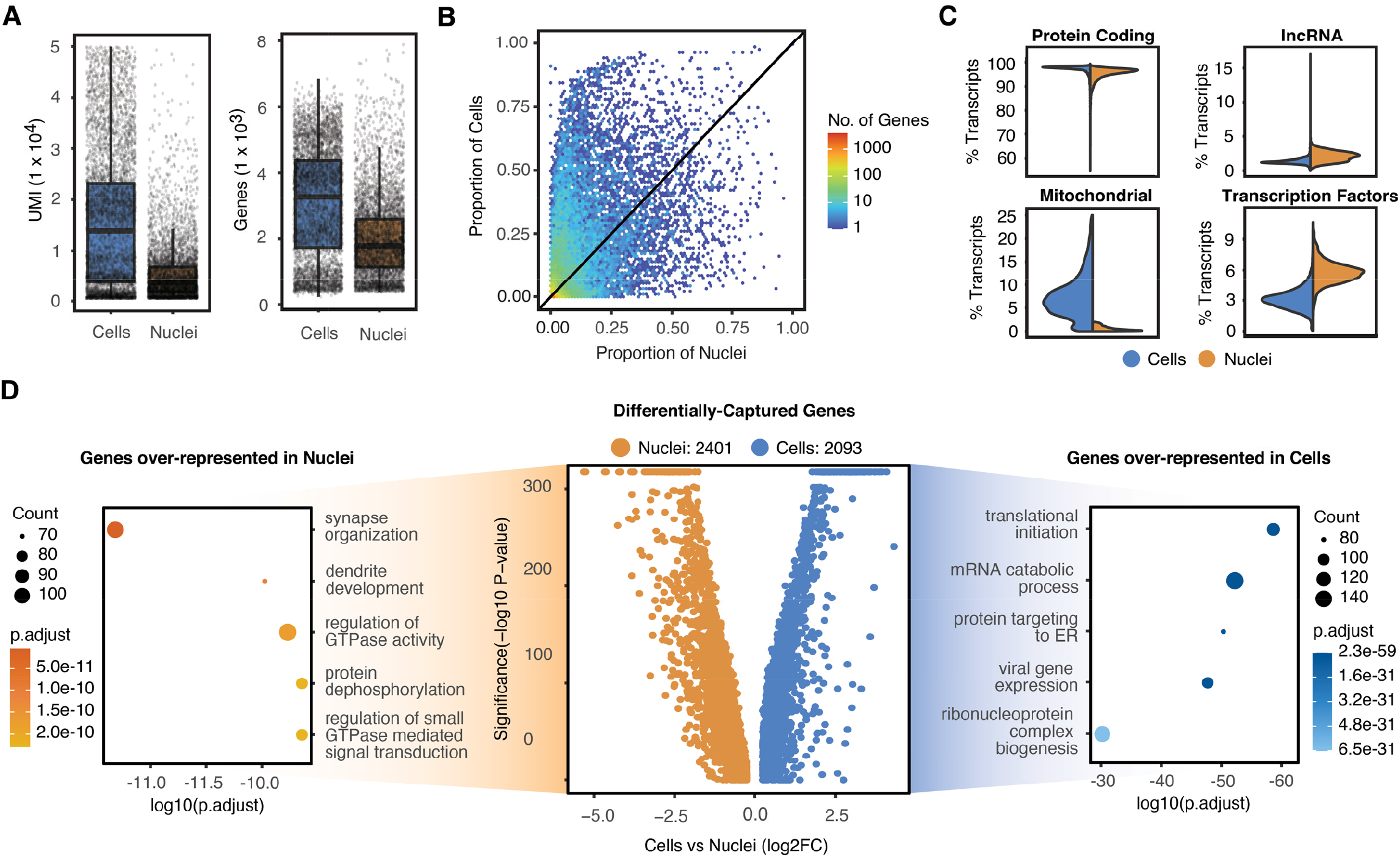
Differences between single cell and single nuclei RNA-sequencing modalities. (A) Boxplots showing UMI capture rate (left) and gene capture rate (right) is higher in cells versus nuclei (B) Binned scatterplot of the proportion of genes detected in cells versus nuclei. (C) Split violin plots displaying the distribution of percentage transcripts that map to protein coding (top left), long non-coding RNA (top right), mitochondrial (bottom left) and transcription factor genes (bottom right). (D) 4494 genes are differentially captured in the combined dataset with 2401 genes expressed higher in nuclei and 2093 expressed higher in cells. The top 5 over-represented gene ontology terms (GO) are displayed on the left (nuclei) and right (cells) of the volcano plot.

By analyzing multiple timepoints of PVR progression, we were also able to investigate the efficiency with which scRNA-Seq and snRNA-Seq captured differences between resting, activated, and fibrotic Müller glial cells. We investigated this by conducting combined UMAP analysis of Müller glia from both scRNA-Seq and snRNA-Seq datasets, incorporating equal number of cells from each dataset generated and timepoint profiled (Fig. 4A). Using well-characterized molecular markers ^35–38^, we then grouped glia into resting, reactive, and fibrotic subgroups (Fig. 4B). Müller glia profiled by scRNA-Seq showed much higher overall enrichment for reactive glia (Fig. 4D,E), while glia profiled by snRNA-Seq were much more likely to be fibrotic (Fig. 4D,E), and somewhat more likely to be in a resting state, even after controlling for injury state. Significant differences in the expression in the relative levels of marker genes of specific glial states are also observed between scRNA-Seq and snRNA-Seq datasets. Pseudotime lineage analysis and RNA velocity analysis of the individual datasets and the combined dataset show overlap of cellular transitions between glial cell states, starting with control cells, followed by reactive and terminating with fibrotic cells (Fig. 4B-D,F, Supplementary Fig. S4A-C). Differential gene expression along pseudotime of the combined dataset also captures the majority of gene changes seen in the individually analyzed datasets (Fig. 4G,H).

**Figure 4:**
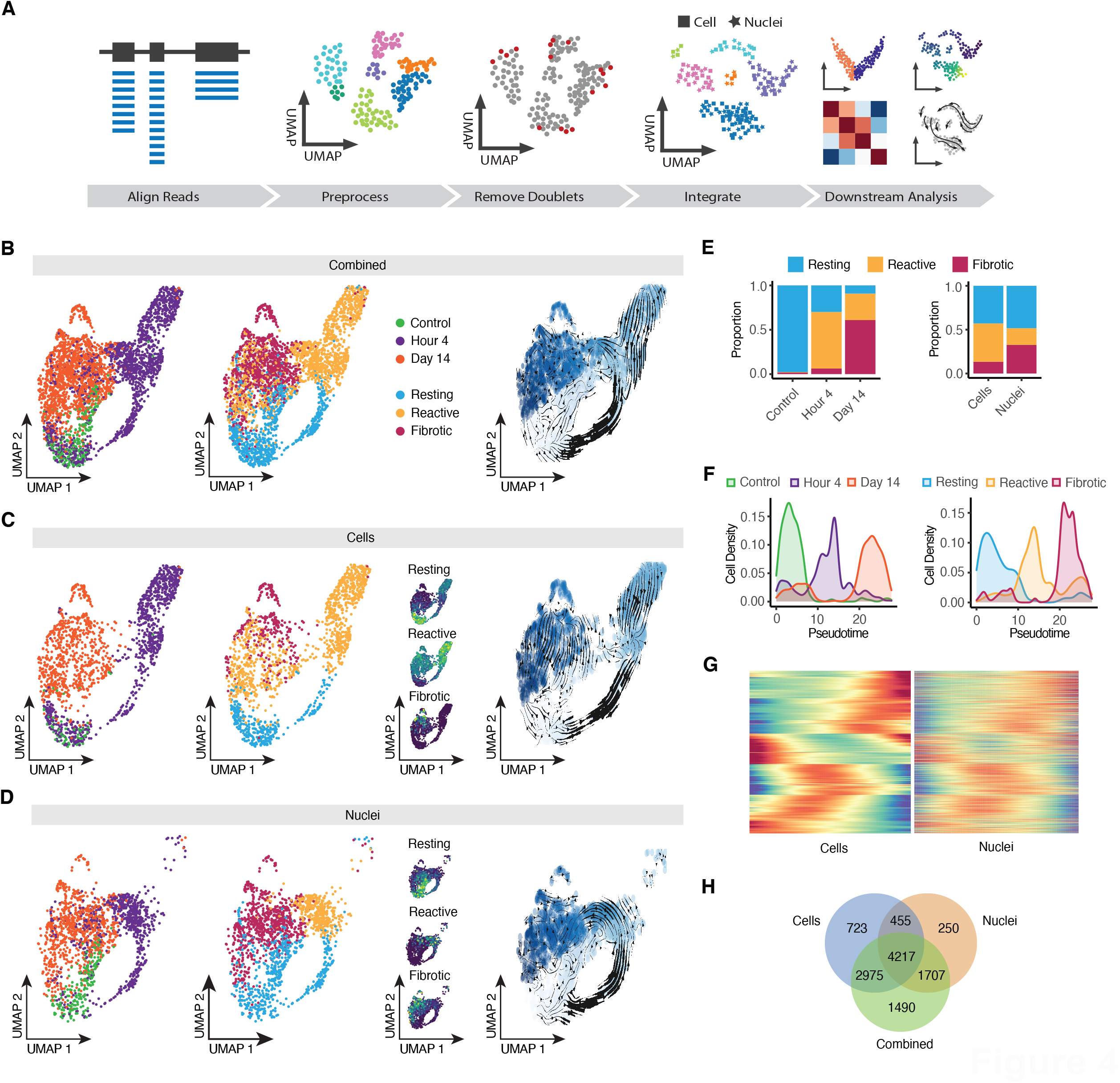
Integration of scRNA and snRNA-seq captures Müller glia transitional states after PVR induction. (A) Schematic summary of integrating and combined analysis of scRNA and snRNA-seq data. (B-D) UMAP embeddings of the combined integrated (B), cells (C) and nuclei (D). Dots are colored by injury timepoints (left), cell state with the respective combined expression of cell state specific genes (middle) and pseudotime with RNA velocity trajectory predictions overlayed. (E) Comparison of Müller glia transitional states proportions between injury timepoints of the combined dataset (left) and sequencing modalities (right). (F) Density plots of injury timepoints (left) or cell states (right) plotted along pseudotime in the combined dataset. (G) Heatmap comparing gene expression dynamics of common highly variable differentially expressed genes along pseudotime between scRNA-seq (left) and snRNA-seq (right). (H) Comparison of combined or modality specific analysis of differentially expressed genes along pseudotime.

## Discussion

In this study, we perform a systematic comparison of scRNA-Seq and snRNA-Seq datasets obtained from a rabbit model of PVR across multiple stages of disease progression. Although both scRNA-Seq and snRNA-Seq capture and accurately profile gene expression in abundant retinal cell types, we observe major differences between these datasets. Overall, cell type proportions captured by snRNA-Seq much more accurately reflect cell composition data obtained from histological analysis. ScRNA-Seq overrepresented Müller glia and microglia, as well as cone photoreceptor and bipolar interneurons, but captures few amacrine cells and hardly any retinal ganglion cells, oligodendrocytes, and astrocytes. This may reflect relative differences in viability during the dissociation and microfluidic steps of library construction and may be partially overcome through the use of rapid methanol fixation or similar approaches ^39^. Furthermore, despite the greater overall efficiency of Müller glia capture with scRNA-Seq, snRNA-Seq more efficiently captures fibrotic glia. Improved recovery of fibrotic cells with snRNA-Seq has previously been reported in kidney ^26^, and may reflect greater difficulty in removing extracellular matrix associated with these cells and thereby obtaining clean dissociation. The greater level of rod and cone photoreceptor contamination seen in scRNA-Seq datasets likely reflects inclusion of mRNA-containing inner segment fragments, which may be less likely to contaminate nuclear preparations, while the greater levels of inner retinal cell contamination seen in snRNA-Seq may simply reflect the overall more efficient recovery of these cell types.

Although these findings generally weigh in favor of routine use of snRNA-Seq for these studies, scRNA-Seq offers several potential overall advantages. The number of transcripts and genes detected in each cellular profile are, as expected, considerably higher than observed with snRNA-Seq. We likewise observe many differences in the expression of individual genes between cell-specific scRNA-Seq and snRNA-Seq profiles. Transcripts encoding certain functional categories of genes are consistently enriched in either scRNA-Seq or snRNA-Seq datasets both across all cell types and in individual cell types. Likewise, several well-characterized marker genes for different Muller glia activation states show dramatic differences in expression levels between scRNA-Seq and snRNA-Seq data obtained from identical biological samples. If resources allow, combined use of both approaches may thus provide a more comprehensive and useful dataset for analyzing progression of retinal disease.

## Methods

### PVR Induction

All animal experiments were performed in accordance with the guidelines for the Use of Animals in Ophthalmic and Vision Research of the Association for Research in Vision and Ophthalmology (ARVO), approved by the Johns Hopkins University’s Institutional Animal Care and Use Committee and in adherence with the ARRIVE 2.0 Guidelines. Dutch Belted rabbits (1.0-4.0kg) obtained from RSI (Robinson Services Inc, Mocksville, NC) were maintained in a temperature-controlled, 12h light cycle environment with *ad libitum* food and water. Briefly, animals were anesthetized with ketamine (35mg/kg IM) and xylazine (5mg/kg IM), intubated, and maintained on isoflurane. Unilateral induction of PVR lesions was achieved via lensectomy, pars plana vitrectomy, and retinotomy followed by retinal cryotherapy and intravitreal autologous platelet-rich plasma injection; contralateral eyes served as controls.

### Cell dissociation

Rabbits were euthanized, and eyes were enucleated and placed in HBSS. Control tissue or PVR lesions were dissected en bloc and transferred to ice cold media containing Hibernate A, 1X B27 supplement and 1X GlutaMAX (ThermoFisher, Waltham, MA). The tissue was then dissociated using the Papain Dissociation System (Worthington, Lakewood, NJ). Briefly, samples were incubated in the papain/DNAse mixture for 30 minutes at 37°C with gentle inversion every 5 minutes until most of the tissue broke apart. The cell suspension was triturated 10 times using a wide bore pipette before being mixed in a 1:1 ratio with 0.1X albumin-ovomucoid inhibitor and DNAse in EBSS. The cell:inhibitor mixture was carefully layered on top of 1X albumin-ovomucoid inhibitor in EBSS and centrifuged at 70 x g for 6 minutes at room temperature. The cell pellet was resuspended in ice cold PBS containing 0.04% bovine serum albumin (BSA) and 0.5 U/μl RNasin Plus RNase inhibitor (Promega, Madison, WI) to achieve a concentration of ∼750-1200 cells/μl. Cell concentration and viability was assessed using a combination of trypan blue and DAPI staining.

### Nuclei lysis

Nuclei were isolated following a modified version of 10x Genomics’ isolation of nuclei from embryonic mouse brain tissue. Harvested PVR lesions were first homogenized in 1 ml of chilled lysis buffer (10 mMTris-HCl, 10 mM NaCl, 3 mM MgCl2, 0.01% Nonidet P40) using an RNase free pestle. This suspension was incubated on ice for 15 minutes and then triturated 10 times using a wide bore pipette. The sample was passed through a 70 μm Flowmi filter and centrifuged at 600 x g for 5 minutes at 4°C. The supernatant was discarded, and the nuclei pellet was resuspended in 1 ml nuclei wash and resuspension buffer (1X PBS with 1% BSA and 0.2 U/μl RNase inhibitor) and filtered through a 40 μm Flowmi filter to remove cell debris and large clumps. The centrifugation, resuspension and filtering was repeated once more before the nuclei were resuspended in the proper volume of nuclei wash and resuspension buffer to achieve a concentration of ∼750-1200 nuclei/μl. Nuclei concentration was assessed using a combination of trypan blue and DAPI staining.

### Single Cell RNA-seq library construction and sequencing

Single cell RNA-seq (scRNA-seq) and single nuclei RNA-seq (snRNA-seq) was performed on dissociated retinal cells or nuclei using the Chromium Next GEM Single Cell 3 ′ Reagent Kits v3.1 (10 ×Genomics, Pleasanton, CA). Briefly, retinal cells or nuclei (∼16,000 cells per sample) were loaded into the 10 × Chromium controller and downstream scRNA-seq or snRNA-seq libraries were generated and indexed by following the manufacturer’s instructions. Libraries were pooled and sequenced on Illumina NovaSeq 6000 targeting 50,000 reads per cell.

### scRNA-seq analysis

Sequencing reads were demultiplexed and aligned to the OryCun2.0 rabbit reference genome using the Cell Ranger 6.1.2 *mkfastq* and *count* pipeline (10x Genomics, Pleasanton, CA), using default parameters for scRNA-seq or using the *--include-introns* flag for snRNA-seq. The generated cell-by-gene count matrix was then used as the input for downstream analysis. The count matrix was analyzed using the Seurat 4.0 R package ^40^. Metadata corresponding to sample, injury time point, and modality was added and merged into cell only, nuclei only or combined Seurat objects. Cells and nuclei that had less than 500 UMIs or greater than 50000 UMI along with cells that had more than 25% mitochondrial content or nuclei that had more than 2% mitochondrial content were filtered out. Doublets were identified and removed using the scDblFinder R package ^41^.

The cell, nuclei and combined datasets underwent normalization, variable feature selection and scaling using the default parameters in Seurat’s *NormalizeData, FindVariableFeatures* and *ScaleData* functions. Principal components were then calculated based on the top 2000 variable features and batch corrected using the Harmony R package ^42^. UMAP dimension reduction was performed on the top 10 corrected principal components and clusters were computed using Seurat’s *FindNeighbors* and *FindClusters* functions. Cell types were then identified in the cell and nuclei datasets using a list of known marker genes that were used previously and then transferred into the combined datasets ^13,35,43^. Cell type similarities between the cell and nuclei datasets were then calculated on all variable features using the MetaNeighbor R package ^44^.

Before identifying differentially expressed genes between the cells and nuclei, the combined dataset was randomly downsampled so that each cell type had equal numbers from either sequencing modality to reduce bias due to over representation of certain cell types in either dataset. Differentially expressed genes between cells and nuclei were determined using the Wilcoxon rank sum test. The clusterProfiler R package was used to perform gene ontology enrichment analysis of biological processes found in the differentially expressed gene lists ^45^.

To determine the different cell states of Müller glia, the Müller glia from the combined dataset was subsetted and randomly downsampled so that each sequencing modality had equal contribution. The data was re-normalized and scaled with the gene number and UMI variables regressed out. Principal components were calculated based on the top 2000 variable features and Harmony batch corrected using the sample ID and modality variables to remove. The cell state scores of Müller glia were calculated by the UCell R package using known genes of resting and reactive glia and fibrotic cells identified in previous studies ^35,46,47^. Using the control cells as the root, the Slingshot R package was used to order the cells along a pseudotime lineage and a negative binomial generalized additive model was fitted using the *fitGAM* function of the tradeSeq R package ^48,49^. An association test was used to determine gene expression changing along the pseudotime lineage. The pseudotime analysis was also replicated and compared using the monocle3 R package and differentially expressed genes along pseudotime were grouped into modules using the *find_gene_modules* function ^50^. RNA velocity was used to infer transcriptional dynamics and predict the future state/transition of individual cells/nuclei in the dataset using the spliced and unspliced counts data generated by velocyto python package and velocity calculated and visualized using the scVelo python package ^51,52^.

## Data and code availability

All rabbit scRNA-seq and snRNA-seq data can be accessed at GEO accession numbers GSE217333. Code used to analyze the datasets in this study can be found at https://github.com/csanti88/pvr_singlecell_vs_singlenucleiRNAseq_2022.

## Acknowledgments

The authors would like to thank the Johns Hopkins Transcriptomics and Deep Sequencing Core for use of the 10x Genomics Chromium controller and library sequencing and all the members of the Singh and Blackshaw labs for comments on the manuscript. This project was funded by Bayer AG as part of a sponsored research collaboration with the Wilmer Eye Institute, Johns Hopkins Hospital to MSS and a Visual Sciences Training grant 2T32EY007143 to CPS.

## Figure legends

**Supplementary Figure S1:**
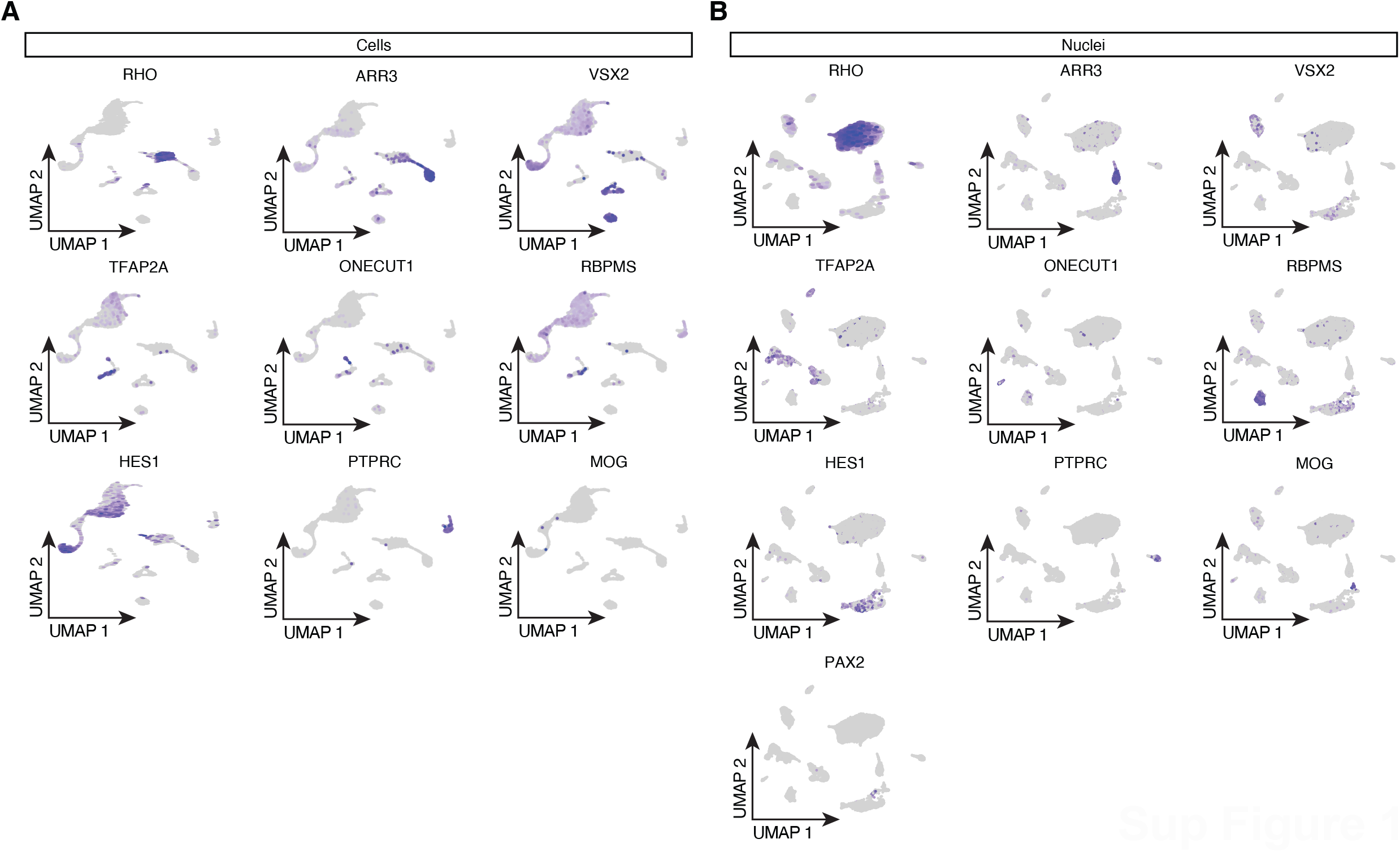
Feature plots of marker gene expression in (A) scRNA-seq and (B) snRNA-seq datasets.

**Supplementary Figure S2:**
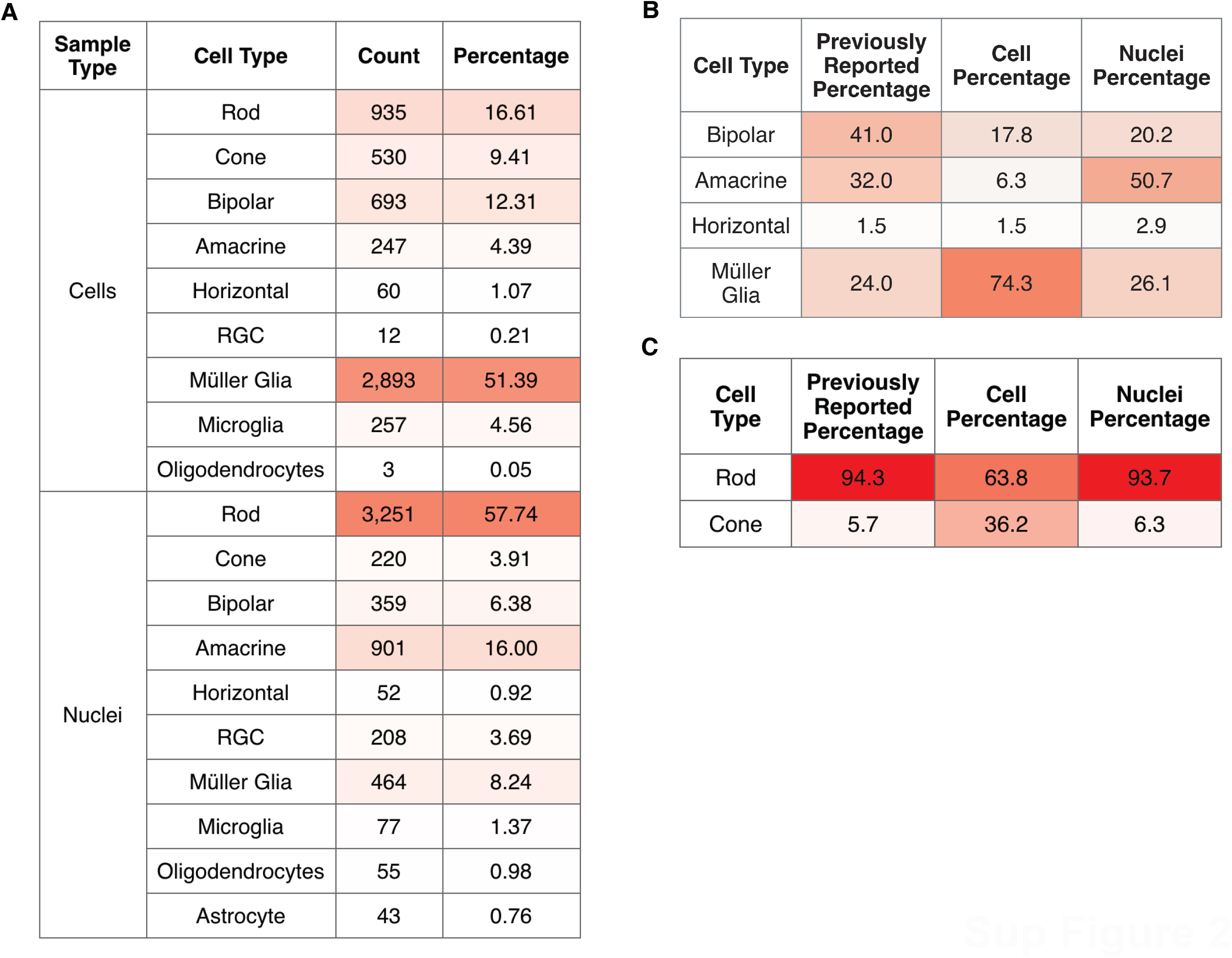
Cell type proportions of the current dataset. (A) Counts and cell type proportion of in the scRNA-seq and snRNA-seq data. (B-C) Comparison of the observed cell and nuclei cell type percentage to previous histological studies for cell types found in the rabbit (B) inner nuclear layer and (C) outer nuclear layer.

**Supplementary Figure S3:**
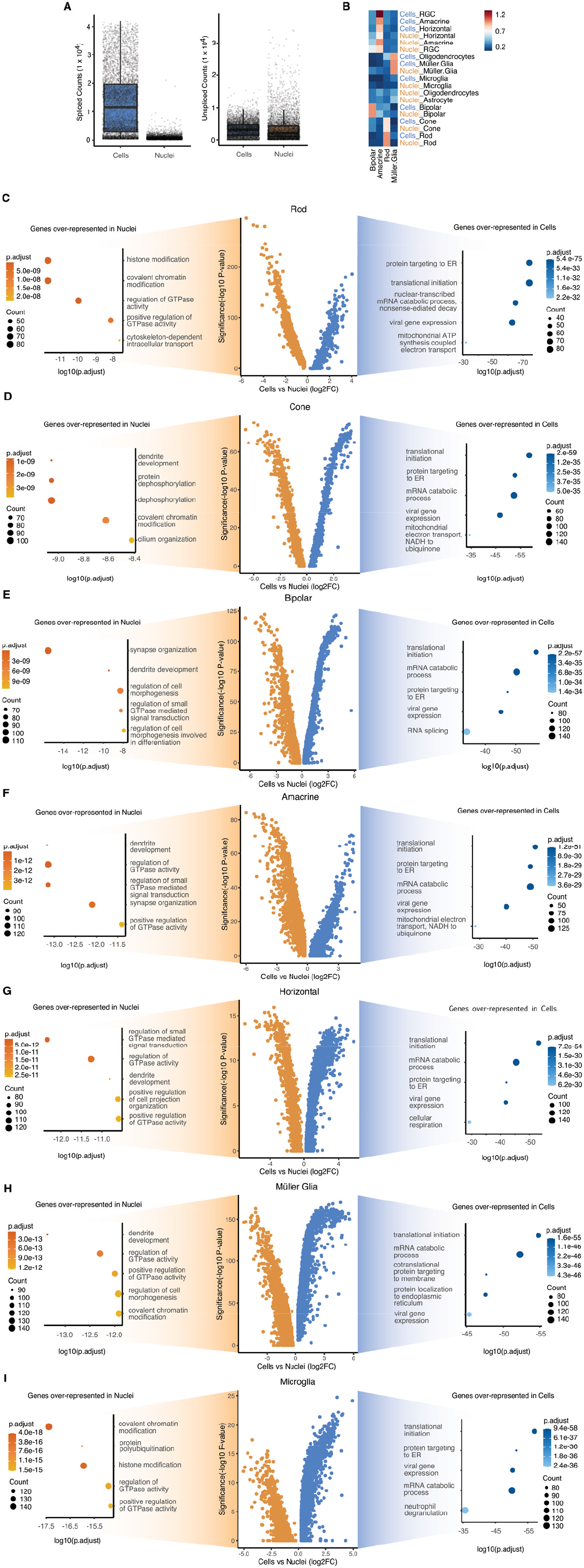
Differences between single cell and single nuclei RNA-sequencing modalities for each cell type. (A) Boxplots showing spliced (left) and unspliced RNA counts between scRNA and snRNA-seq. (B) Expression levels of enriched bipolar, amacrine, rod and Müller glia genes in each cell type. (C-I) Differentially captured genes and top 5 over-represented GO terms between scRNA and snRNA-seq for (C) rod, (D) cone, (E) bipolar, (F) amacrine, (G) horizontal, (H) Müller glia and (I) microglial cell types.

**Supplementary Figure S4:**
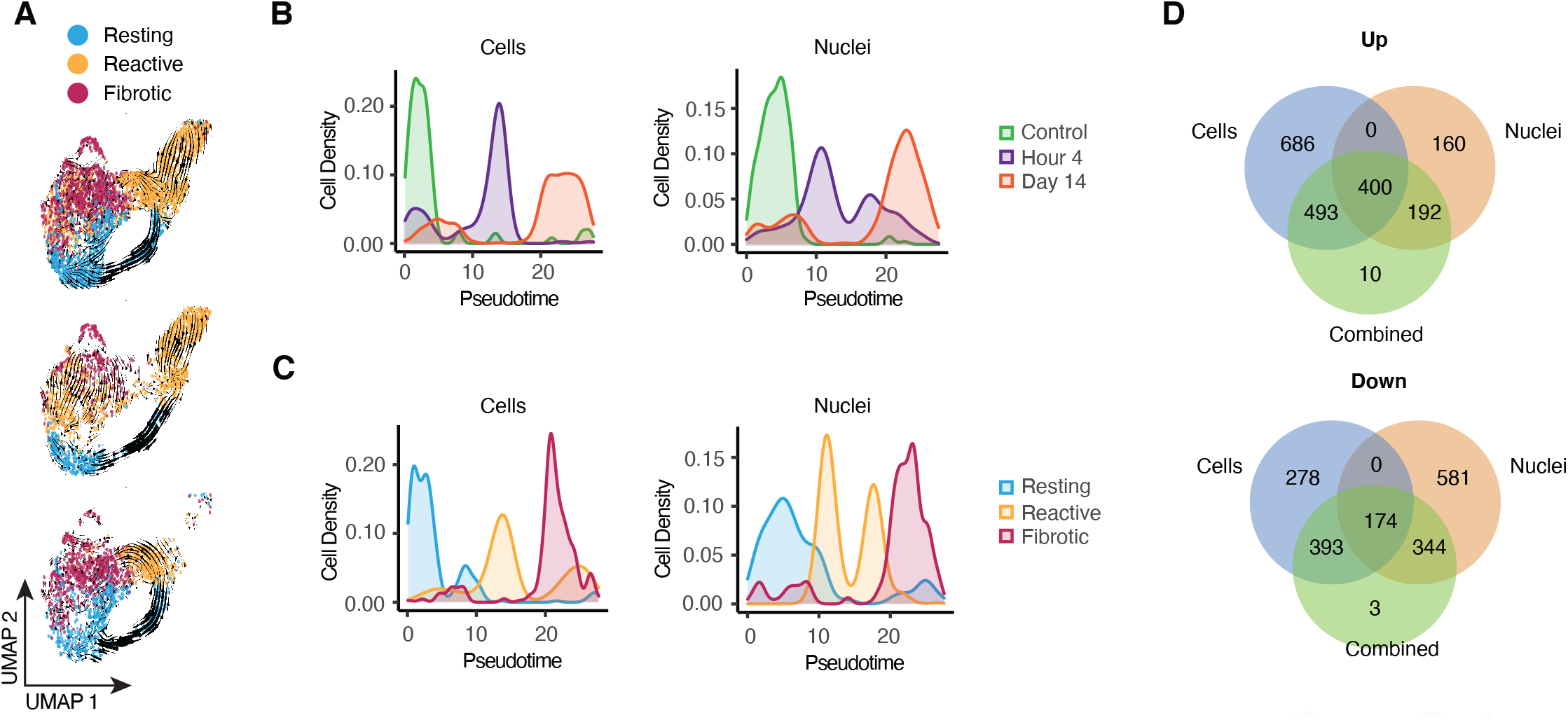
Combined analysis of injured Müller glia from the scRNA and snRNA-seq datasets. (A) RNA velocity trajectory predictions overlayed on Muller glia from the combined (top), cells (middle) or bottom (nuclei) datasets, with each cell colored by cell state. (B-C) Density plots of injury timepoint (B) or cell state (C) plotted along pseudotime separated by sequencing modality. (D) Comparison of combined or modality specific analysis of differentially expressed genes between control and injury by Wilcoxon rank sum test on pseudobulk gene profiles separated by genes that are upregulated (top) or downregulated (bottom) after injury.

